# High Efficiency Gene Editing And Phenotype Generation In Avian Embryos Using Lipid Nanoparticles As A Vector Delivery Platform

**DOI:** 10.64898/2026.10.13.753869

**Authors:** Juan P. Zanin, Benjamin M.J Collerton, Samatha K. Nguyen, William N. Feist, Melissa L Coquelin, Jeffrey N. Stout, Bailey B. Blair, James B. Papizan, Xiaoming Zhan, Richard A Schneider, Anna L. Keyte

## Abstract

Birds are a longstanding developmental biology model because easy access to their fertilized eggs allows for direct observation and experimental manipulation during any stage of embryonic growth prior to hatching. However, functional genetics studies in avian embryos continue to rely primarily on in ovo electroporation and viral transduction, each of which is limited by variable efficiency, procedure-induced embryonic defects, substantial embryo mortality, and ineffective delivery of gene editing reagents such as CRISPR/Cas9. Here we establish lipid nanoparticle (LNP)-mediated delivery as a reproducible alternative for manipulating gene expression in avian embryos with high viability. Using the chicken embryo (*Gallus gallus*), we targeted migrating cranial NCCs, which contribute extensively to the craniofacial skeleton. We optimized LNP formulation and injection timing and frequency to build a robust and adaptable pipeline that offers efficient delivery of plasmid DNA and RNA cargos. We show that LNP delivery can support the piggyBac transposon system, enabling persistent reporter expression and long-term labeling of cranial NCC derivatives. To test if LNP-mediated delivery of RNA-based CRISPR/Cas9 components can achieve efficient somatic genome editing in ovo, we targeted Runt-related transcription factor 2 (*RUNX2)*, which is required for osteogenesis. Embryos with *RUNX2* edits in their cranial NCCs show a severe loss and disruption of craniofacial ossification on the targeted side of their craniofacial skeletons, demonstrating the potential of this approach for rapid functional interrogation of gene function in vivo. We also show that LNPs can mediate transfection in domestic pigeon (*Columba livia*) and Pekin duck (*Anas platyrhynchos)* embryos, indicating this method can be deployed across avians as a vector delivery platform for studying development, disease, and evolution.

## INTRODUCTION

Avian embryos have long served as powerful models for vertebrate developmental biology because of their accessibility, experimental tractability, and amenability to direct manipulation in ovo. The chicken (*Gallus gallus*) embryo, in particular, has been widely used to investigate tissue patterning, organogenesis, and craniofacial development, while the diversity of avian species provides a valuable comparative framework for studying the developmental mechanisms underlying morphological evolution (Schneider and Helms, 2003; Hall et al., 2014; Ealba et al., 2015; Schneider, 2024). Despite these advantages, efficient genetic manipulation of avian embryos remains challenging. Electroporation is widely used for nucleic acid delivery and has enabled gain- and loss-of-function studies, including CRISPR/Cas9-mediated somatic genome editing (Véron et al., 2015), but its efficiency depends strongly on electrode orientation, developmental stage, and the accessibility and geometry of the target tissue (Simkin et al., 2014). Application of electrical pulses can also perturb embryonic tissues and compromise viability. Viral vectors such as RCAS and RCASBP provide an alternative for sustained transgene expression but are constrained by limited cargo capacity, progressive viral spread that can reduce spatial and temporal control, and differences in susceptibility among avian strains and species (Hughes, 2004). These limitations are particularly relevant when targeting migratory, spatially dispersed, or otherwise difficult-to-access cell populations. Genome-editing approaches present an additional challenge because they require efficient intracellular delivery of multiple components and may benefit from repeated administration to increase cellular coverage. Thus, a complementary platform capable of reproducible, spatially localized, and repeated delivery of diverse nucleic acid cargos would substantially expand the experimental toolkit available for avian systems.

Lipid nanoparticles (LNPs) have emerged as highly effective non-viral vehicles for nucleic acid delivery. Modern LNP formulations protect nucleic acid cargo from degradation, promote cellular uptake, and facilitate cytoplasmic release, enabling potent delivery of messenger RNA in vivo (Hou et al., 2021). LNP composition, particularly the identity of the ionizable lipid, strongly influences delivery efficiency (Hou et al., 2021). Unlike plasmid-based expression, direct mRNA delivery allows for rapid but transient production of the encoded protein without requiring nuclear entry or integration of an expression vector. Importantly, LNPs can also deliver genome-editing machinery: co-delivery of Cas9 mRNA and guide RNA has produced efficient somatic genome editing in mammalian tissues, demonstrating that transient delivery of editing components can generate permanent genomic modifications (Finn et al., 2018; Gillmore et al., 2021). Together with their ability to accommodate different classes of nucleic acid cargo, these properties make LNPs an attractive platform for experimental manipulation of developing tissues.

LNP delivery may be particularly well suited to developmental systems because local administration can spatially restrict exposure to a defined embryonic region without application of an electrical field, while repeated injections can target dynamic cell populations at successive developmental time points. Cranial neural crest cells (NCCs) provide a useful model for evaluating these capabilities. During early development, cranial NCCs continuously migrate from the neural tube into and through the cephalic mesenchyme before contributing extensively to the craniofacial skeleton (Noden and Schneider, 2006). Repeated administration could therefore access newly arriving NCCs that were unavailable during earlier treatment while also re-exposing previously targeted cells to additional cargo. This combination of spatial control, repeat dosing, and cargo flexibility could facilitate transient gene expression, persistent genetic labeling, and genome editing within a common delivery framework. Despite the general development of LNP technology for therapeutic applications, its potential as a general experimental platform for genetic manipulation of developing avian embryos in ovo remains largely unexplored.

Here, we establish LNP-mediated nucleic acid delivery as a versatile platform for genetic manipulation of avian embryos in ovo. Using migrating cranial NCCs as a challenging developmental target, we evaluate LNP formulations for localized mRNA delivery and determine whether repeated administration can increase cellular coverage and permit sequential cargo delivery. We further test the versatility of the platform for persistent genetic labeling using the piggyBac transposon system, which promotes genome integration and stable long-term transgene expression. Specifically, we test the applicability of this system in domestic pigeon (*Columba livia*) and Pekin duck (*Anas platyrhynchos)* embryos, and determine whether LNPs can deliver CRISPR/Cas9 components at sufficient efficiency to generate a developmental phenotype in vivo. Together, these experiments establish a common LNP-based framework for manipulating gene expression, achieving persistent labeling of lineages and cells, and performing functional genome editing in developing avian embryos.

## RESULTS

### Ionizable lipid screening demonstrates efficient LNP-mediated mRNA delivery in chicken fibroblasts

To identify LNP formulations suitable for avian applications, we first compared four well-characterized ionizable lipid formulations for delivery of mCherry mRNA to cultured chicken fibroblasts. Untreated control cells showed negligible fluorescence, whereas all four LNP formulations produced detectable reporter expression (Fig. 1A–E). SM102, DLin-MC3-DMA (MC3), and DLin-KC2-DMA (KC2) transfected more than 90% of cells, whereas LP-01 transfected approximately 55% (Fig. 1F). Mean fluorescence intensity further distinguished the formulations: MC3 and KC2 produced the strongest reporter signal, while SM102 and LP-01 were lower (Fig. 1G). These results established that LNPs can efficiently deliver exogenous mRNA to chicken cells and provided a basis for testing the platform in ovo.

**Fig. 1.**
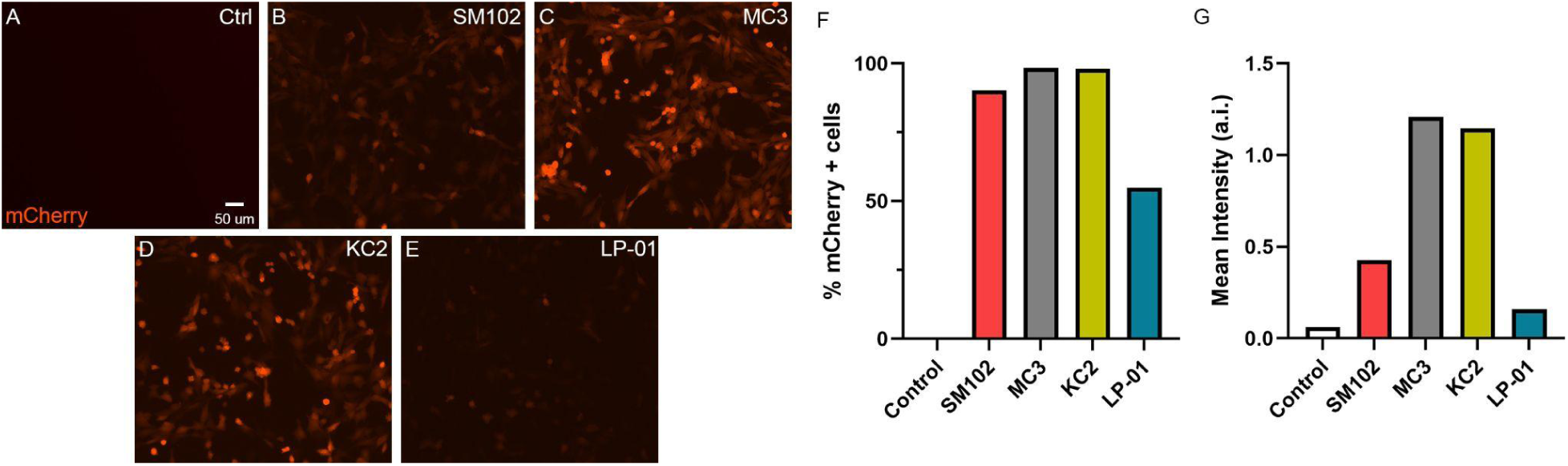
Ionizable lipid screening demonstrates efficient LNP-mediated mRNA delivery in chicken fibroblasts. Chicken fibroblasts were transfected with lipid nanoparticles (LNPs) encapsulating mCherry mRNA and formulated with different ionizable lipids. (A–E) Representative fluorescence images of untreated control cells (A) and chicken fibroblasts treated with LNPs formulated with SM102 (B), MC3 (C), KC2 (D), or LP-01 (E). (F) Quantification of transfection efficiency, expressed as the percentage of mCherry-positive cells for each LNP formulation and untreated control. (G) Quantification of mCherry mean fluorescence intensity (MFI) following treatment with the indicated LNP formulations.

### LNP-mediated mRNA delivery enables spatially restricted transfection of the developing chicken embryo in ovo

We next asked whether LNPs could mediate localized mRNA delivery in the developing chicken embryo. Hamburger-Hamilton (HH) 9/10 stage embryos were injected unilaterally with GFP mRNA-containing SM102 LNPs into the right cephalic mesenchyme adjacent to the neural tube, targeting migrating cranial NCCs, and analyzed 24 hours later. Uninjected controls showed no GFP fluorescence (Fig. 2A, B), whereas SM102 LNPs produced strong reporter expression confined predominantly to the injected cephalic region (Fig. 2C, D). Higher-magnification analysis demonstrated robust GFP expression on the injected side of the head (Fig. 2E, H), with no detectable fluorescence on the contralateral non-injected side (Fig. 2F, I). Frontal views further demonstrated the unilateral distribution of GFP expression within the targeted cephalic mesenchyme, with no detectable fluorescence in the contralateral side or the neural tube (Fig. 2G, J). Together, these observations show that localized LNP administration can produce spatially restricted mRNA delivery in ovo.

**Fig. 2.**
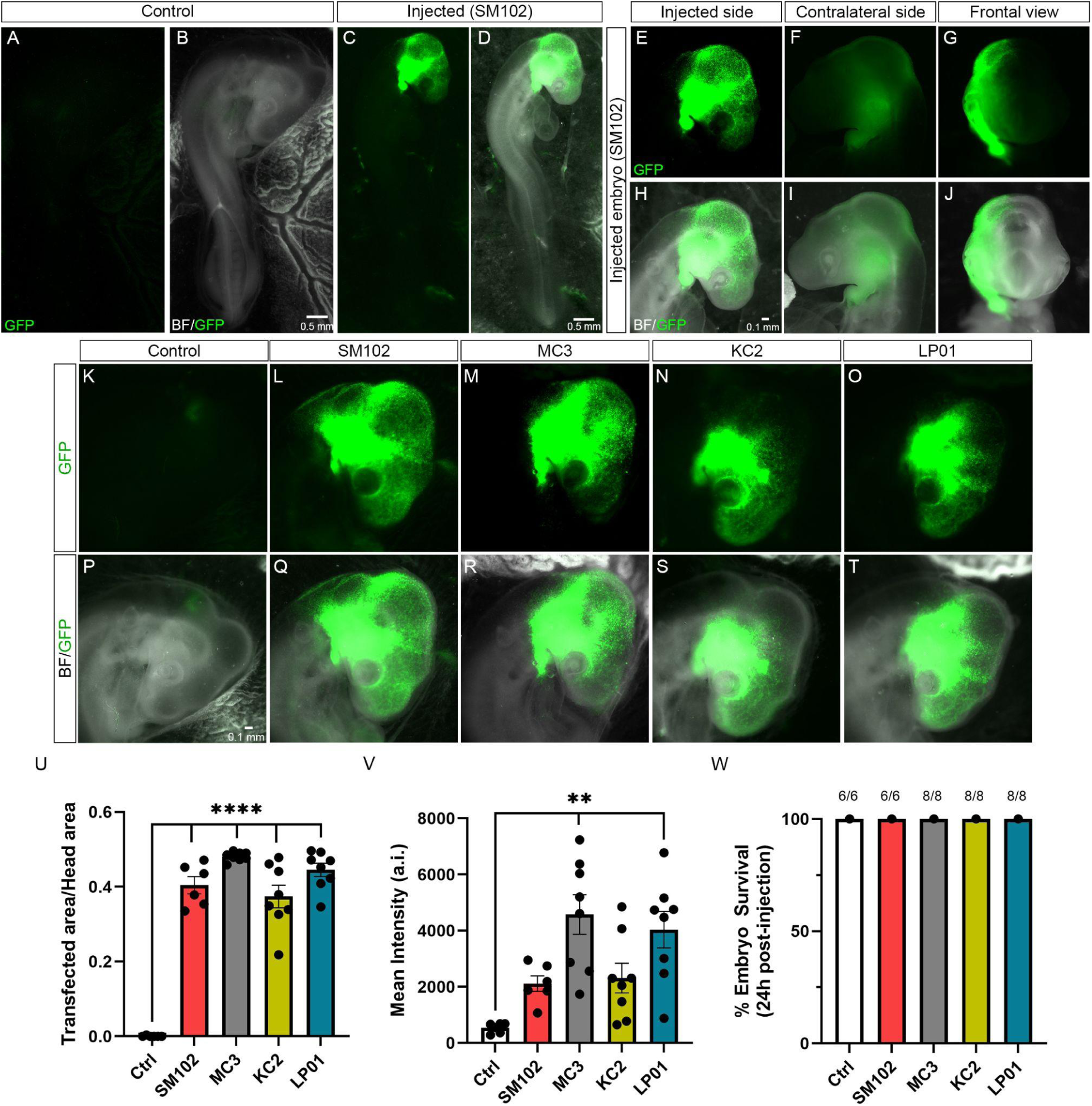
LNP-mediated mRNA delivery enables spatially restricted transfection of the developing chicken embryo in ovo. HH9/10 chicken embryos were injected with LNPs encapsulating GFP mRNA into the right cephalic mesenchyme adjacent to the neural tube and analyzed 24 hours after injection. (A–D) Whole-embryo images showing an uninjected control embryo (A, B) and an embryo injected with SM102-containing LNPs (C, D). GFP fluorescence is shown in A, C, and corresponding bright-field/GFP merged images are shown in B, D. (E–J) Higher-magnification views of the head of an SM102-injected embryo. GFP fluorescence is shown for the injected side (E), contralateral non-injected side (F) and frontal view (G), with corresponding bright-field/GFP merged images shown in H–J, respectively. (K–T) Representative images of the cephalic region following injection of GFP mRNA-containing LNPs formulated with different ionizable lipids. GFP fluorescence images are shown for uninjected control (K), SM102 (L), MC3 (M), KC2 (N) and LP-01 (O), with corresponding bright-field/GFP merged images shown in P–T, respectively. (U) Quantification of GFP-positive area relative to the total head area following treatment with the indicated LNP formulations. (V) Quantification of GFP mean fluorescence intensity within the targeted cephalic region. (W) Embryo survival 24 hours after injection with the indicated LNP formulations. Numbers above bars indicate surviving embryos/total embryos. Data are presented as mean ± s.e.m.; individual points represent independent embryos.

We next compared the four ionizable lipid formulations evaluated in chicken fibroblasts for their ability to deliver GFP mRNA in ovo. GFP images are shown for uninjected controls (Fig. 2K, P) and embryos injected with LNPs formulated with SM102 (Fig. 2L, Q), MC3 (Fig. 2M, R), KC2 (Fig. 2N, S), or LP-01 (Fig. 2O, T). GFP expression was readily detected within the cephalic region following injection with each formulation. Quantification of GFP-positive area relative to the total head area showed reporter expression across at least 40% of the head for all four formulations, compared with negligible signal in uninjected controls (Fig. 2U). The similar spatial coverage observed across formulations further demonstrates the reproducibility of the injection procedure and indicates that the anatomical distribution of transfection is largely determined by the injection site. In contrast, mean GFP fluorescence intensity differed among formulations, with MC3 and LP-01 producing the strongest reporter signal (Fig. 2V), suggesting increased mRNA delivery and/or expression within the targeted tissue. Localized LNP delivery was also well tolerated. All embryos survived to 24 hours after injection across the four formulations tested (Fig. 2W), demonstrating high short-term viability under these conditions.

### Repeated LNP administration increases coverage and enables sequential targeting of migrating cranial neural crest cells

Because cranial NCC migration from the dorsal neural tube into the cephalic mesenchyme is a dynamic process that occurs over multiple hours, we asked whether repeated LP-01 LNP delivery could expand coverage of this cell population. Beginning at HH9/10, embryos received one, two, or three unilateral LP-01 LNP injections into the cephalic mesenchyme. For the single-injection condition, GFP and mCherry mRNAs were co-delivered during the same LNP administration, whereas the two- and three-injection conditions received the reporter cargos in sequential injections separated by 2–3 hours (Fig. 3A). Embryos were collected 24 hours after the first injection, at approximately HH15/16. Repeated administration remained well tolerated, with survival near or above 90% across injection regimens (Fig. 3N).

**Fig. 3.**
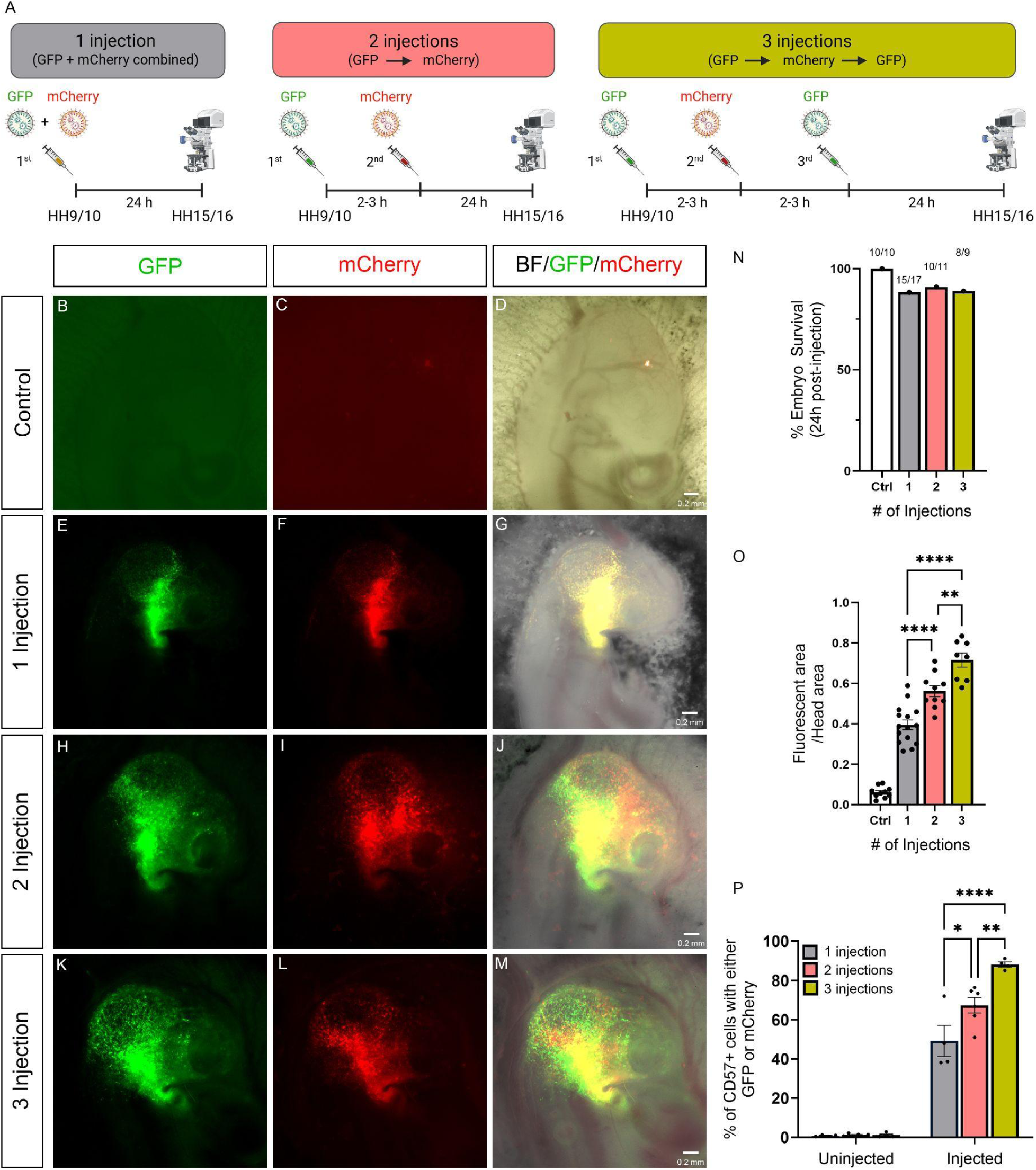
Repeated LP-01 LNP administration increases coverage and enables sequential targeting of migrating cranial NCCs. (A) Schematic of the sequential LP-01 LNP injection strategy. (N) Embryo survival 24 hours after the first injection for uninjected controls and embryos receiving one, two or three LP-01 LNP injections. Numbers above bars indicate surviving embryos/total embryos. For the one-injection group, GFP and mCherry mRNAs were co-delivered during the same injection; for the repeated-injection groups, reporter cargos were delivered sequentially at 2–3 h intervals. (B–D) Uninjected control embryo. Representative embryos following one (E–G), two (H–J) or three (K–M) LNP injections. (O) Quantification of reporter-positive (GFP+ and/or mCherry+) area relative to total head area following one, two or three LNP injections. All injection groups exhibited significantly greater reporter-positive area than uninjected controls (*P*<0.0001); selected comparisons between injection regimens are indicated in the graph. (P) Quantification of reporter-positive (GFP+ and/or mCherry+) cranial NCCs. Cephalic mesenchymal tissue from the injected and contralateral non-injected sides was dissected and dissociated, and CD57/HNK-1-positive NCCs were identified by flow cytometry. Data are presented as mean ± s.e.m.; individual points represent independent embryos. Statistical significance is indicated in the figure; **P<0.01, ****P<0.0001.

Reporter distribution expanded progressively with repeated administration. A single injection produced localized GFP and mCherry expression within the targeted mesenchyme, whereas two and three injections produced increasingly broad reporter coverage (Fig. 3B–M). Merged images contained both overlapping and spatially distinct reporter-positive regions, consistent with sequential injections re-targeting some previously exposed cells while also reaching cells that entered the injection field later. Quantification confirmed this expansion: reporter-positive area increased from approximately 40% after one injection to 55–60% after two and approximately 70% after three (Fig. 3O). All injection regimens resulted in significantly greater reporter-positive area than uninjected controls (*P*<0.0001).

To test whether this increase reflected greater coverage of cranial NCCs specifically, cephalic mesenchyme was dissociated and analyzed by flow cytometry using CD57/HNK-1 to identify a NCC-enriched population. Reporter-positive HNK-1+ cells were negligible on the uninjected side. On the injected side, approximately half of HNK-1+ cells were reporter-positive after one injection, increasing to approximately 65–70% after two injections and nearly 90% after three (Fig. 3P).

These data show that repeated LNP administration can progressively expand delivery across a migrating cell population while also permitting sequential exposure of previously targeted cells. This feature is particularly useful for developmental experiments in which target populations change over time or where efficient genome editing may benefit from repeated or combinatorial cargo delivery.

### LNP-mediated piggyBac delivery supports persistent labeling of cranial neural crest derivatives

Having established efficient transient mRNA delivery using repeated LNP administration, we next asked whether the platform could support persistent genetic labeling using the piggyBac transposon system. HH9/10 chicken embryos received three unilateral injections of LNPs into the cranial mesenchyme adjacent to the neural tube at 2–3 h intervals. LNPs delivered piggyBac transposase together with a transposon donor plasmid encoding EGFP, with the goal of maintaining reporter expression in transfected cells and their descendants.

Repeated LP-01 LNP administration was compatible with continued embryonic development. Approximately 90% of injected embryos survived the first 24 hours (25/28), with 70% survival at 7 days (19/27), 73% at 10 days (11/15), and 50% at 13 days post-injection (6/12) (Fig. 4A). Thus, high acute survival was maintained following repeated administration, and a substantial proportion of injected embryos remained viable through later stages of development.

**Fig. 4.**
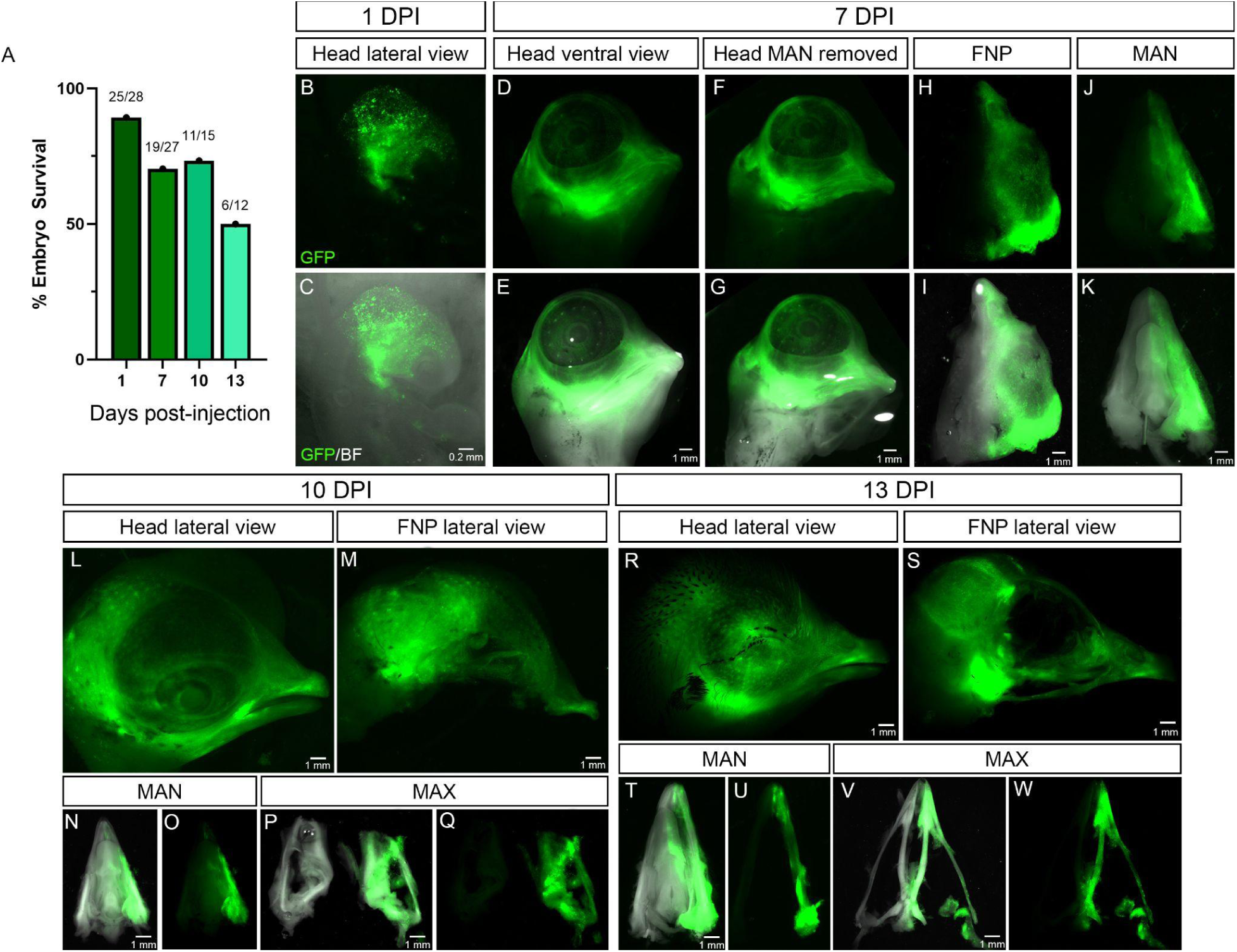
LNP-mediated piggyBac delivery enables persistent labeling of cranial neural crest derivatives throughout craniofacial development. HH9/10 chicken embryos received three unilateral injections of LP-01 LNPs carrying the piggyBac reporter system into the cephalic mesenchyme adjacent to the neural tube, with injections separated by 2–3 h. (A) Embryo survival at 1, 7, 10 and 13 days post-injection. Numbers above bars indicate surviving embryos/total embryos analyzed. (B–W) Representative fluorescence images showing persistent EGFP expression at 1, 7, 10 and 13 days post-injection (DPI). (B, C) Lateral view of the head at 1 DPI showing GFP fluorescence (B) and the corresponding GFP/bright-field merged image (C). (D–K) Craniofacial reporter expression at 7 DPI. Ventral views of the intact head (D, E) and head following removal of the mandible (F,G) are shown, together with dissected frontonasal process (FNP)-derived (H,I) and mandibular (MAN) (J, K) structures. (L–Q) Reporter expression at 10 DPI, including lateral views of the head (L) and FNP-derived region (M) and dissected mandibular (N, O) and maxillary (MAX) (P, Q) structures. (R–W) Persistent reporter expression at 13 DPI, including lateral views of the head (R) and FNP-derived region (S) and dissected mandibular (T, U) and maxillary (V, W) structures. Corresponding GFP and GFP/bright-field images demonstrate persistent reporter expression within cranial NCC-derived tissues throughout craniofacial development. Scale bars are indicated within individual panels. DPI, days post-injection.

EGFP expression was readily detected within the cephalic region at 1 day post-injection (DPI), confirming successful early delivery of the piggyBac reporter system (Fig. 4B, C). Reporter expression persisted as embryos underwent extensive craniofacial growth and morphogenesis. At 7 DPI, ventral views of the head showed widespread EGFP-positive tissue within the developing craniofacial region (Fig. 4D, E), which remained evident following removal of the mandible (Fig. 4F, G). Dissection of individual craniofacial structures further revealed persistent reporter expression within frontonasal process (FNP)-derived tissues (Fig. 4H, I) and the mandible (Fig. 4J, K). Persistent labeling was also evident at later time points. At 10 DPI, lateral views of the head showed extensive EGFP expression throughout the craniofacial region (Fig. 4L), including the frontonasal (FNP)-derived region (Fig. 4M). Dissected mandibular (Fig. 4N, O) and maxillary (Fig. 4P, Q) structures retained strong reporter expression, demonstrating persistence of the labeled population within differentiated craniofacial derivatives. By 13 DPI, GFP-positive tissue remained readily detectable in lateral views of the head and FNP-derived region (Fig. 4R, S), as well as within dissected mandibular (Fig. 4T, U) and maxillary (Fig. 4V, W) skeletal elements.

Because LNPs were administered adjacent to the neural tube at HH9/10, before cranial NCCs had completed colonization of the facial prominences, the persistence and distribution of EGFP-positive cells across FNP-, mandibular- and maxillary-derived structures are consistent with early targeting of migrating cranial NCCs followed by maintenance of reporter expression in their descendants. Reporter expression persisted for at least 13 days despite extensive tissue growth, differentiation and morphogenetic remodeling. Thus, LNP-mediated delivery can support not only transient mRNA expression but also persistent piggyBac-mediated genetic labeling, substantially expanding the range of experimental applications accessible through the same in ovo delivery platform.

### LNP-mediated piggyBac delivery enables persistent genetic labeling across avian species

To determine whether the formulations we derived for LNP-mediated delivery could be extended beyond chicken to other avian species, we tested the system in domestic pigeon (*Columba livia*) and duck (*A. platyrhynchos*) embryos. Embryos were injected at HH9/10 with LP-01 LNPs carrying the piggyBac reporter system into the cephalic mesenchyme adjacent to the neural tube, targeting migrating cranial NCCs. Robust GFP expression was detected within the targeted cephalic region 1 day post-injection (DPI) in both pigeon (Fig. 5A,B) and duck (Fig. 5M,N) embryos, demonstrating efficient early delivery of the reporter system across distinct avian species.

**Fig. 5.**
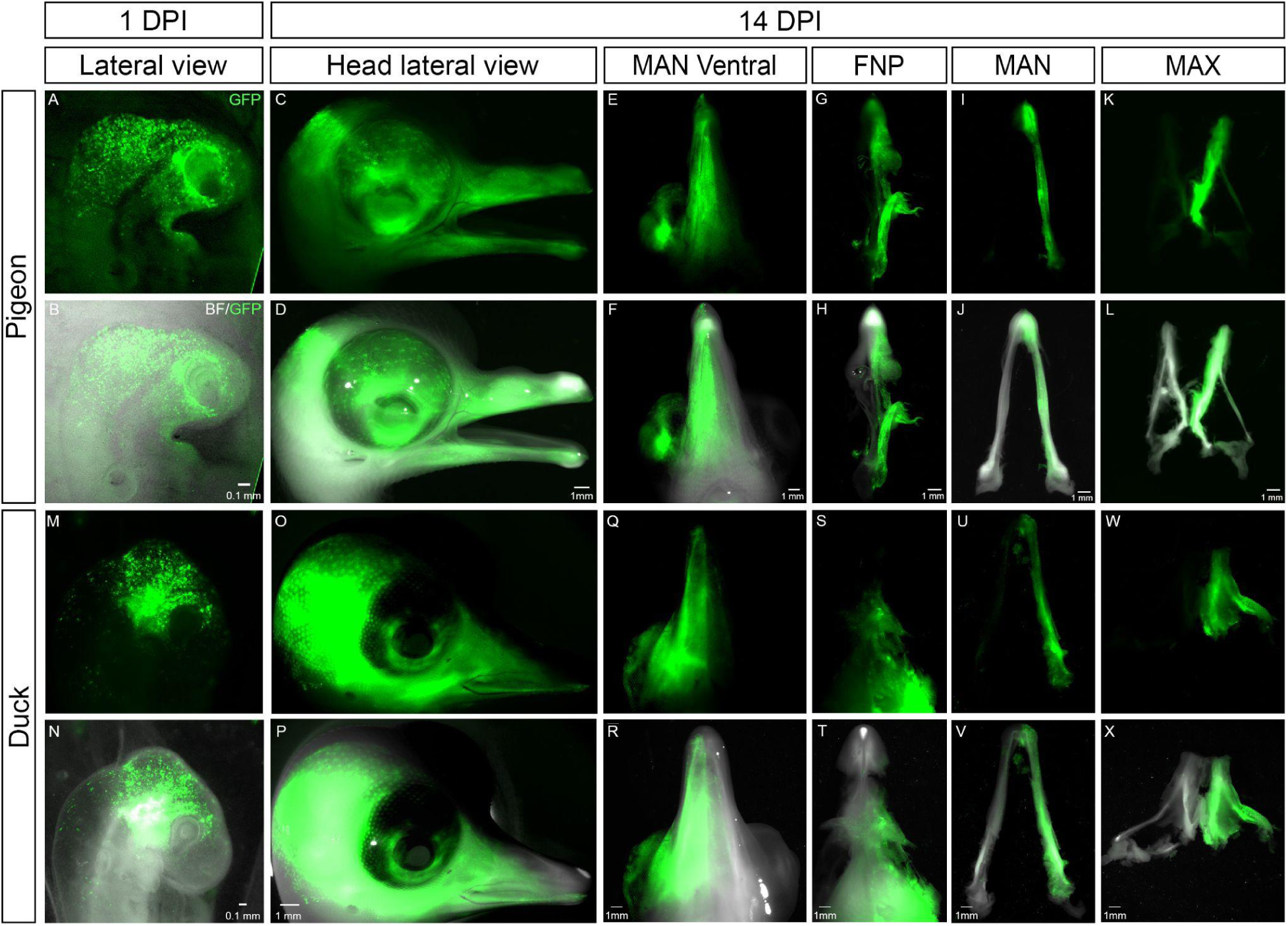
LNP-mediated piggyBac delivery enables persistent genetic labeling of cranial neural crest derivatives across avian species. Pigeon (*C. livia*) and duck (*A. platyrhynchos*) embryos were injected at HH9/10 with LP-01 LNPs carrying a piggyBac-based reporter system into the cephalic mesenchyme adjacent to the neural tube. (A–L) LNP-mediated delivery in pigeon embryos. Representative embryo at 1 day post-injection (DPI), showing GFP fluorescence (A) and corresponding bright-field/GFP merged image (B). At 14 DPI, persistent reporter expression is shown in lateral views of the head (C, D), ventral views of the mandibular region (E, F), dissected frontonasal process (FNP)-derived structures (G, H), dissected mandibular skeletal elements (I, J) and maxillary derivatives (K, L). (M–X) Corresponding analysis in duck embryos. GFP fluorescence and bright-field/GFP merged images are shown at 1 DPI (M, N) and 14 DPI for lateral views of the head (O, P), ventral views of the mandibular region (Q, R), FNP-derived structures (S, T), dissected mandibular skeletal elements (U, V) and maxillary derivatives (W, X). Persistent GFP expression within corresponding craniofacial structures in pigeon and duck demonstrates long-term labeling of cranial NCC derivatives following LNP-mediated delivery of the piggyBac system and supports the applicability of this delivery strategy across avian species. Scale bars are indicated within individual panels. DPI, days post-injection.

We next determined whether reporter expression persisted as the targeted NCC populations contributed to later craniofacial structures. At 14 DPI, strong GFP expression remained readily detectable throughout the craniofacial region in both pigeon (Fig. 5C,D) and duck (Fig. 5O,P) embryos. In both species, reporter expression remained spatially associated with the targeted craniofacial tissues despite extensive growth and morphogenesis, consistent with persistent labeling following LNP-mediated delivery of the piggyBac system.

Examination of individual craniofacial structures further demonstrated comparable targeting of NCC derivatives across species. Ventral views of the mandible showed extensive GFP-positive tissue in pigeon (Fig. 5E,F) and duck (Fig. 5Q,R). Dissected FNP-derived structures similarly retained strong reporter expression in both species (pigeon, Fig. 5G,H; duck, Fig. 5S,T), as did dissected mandibular skeletal elements (pigeon, Fig. 5I,J; duck, Fig. 5U,V) and maxillary derivatives (pigeon, Fig. 5K,L; duck, Fig. 5W,X). Thus, cranial NCC targeted during early migration retained reporter expression as they contributed to differentiated derivatives of the frontonasal, mandibular and maxillary prominences.

Together with the chicken experiments, these results demonstrate successful LNP-mediated genetic delivery in representative species from three highly divergent avian orders (i.e., Galliformes, Columbiformes, and Anseriformes) and show that the same localized delivery strategy can target comparable craniofacial populations. The reproducibility of reporter distribution across species with substantially different craniofacial morphologies supports the applicability of LNPs as a genetic delivery platform for avian embryos more broadly. Combined with persistent piggyBac-mediated labeling, this cross-species portability provides a framework for comparative functional experiments aimed at testing how genetic and regulatory mechanisms contribute to species-specific development and morphological evolution.

### LNP-mediated CRISPR/Cas9 delivery enables efficient *RUNX2* editing and produces craniofacial skeletal defects

Having established LNP-mediated delivery as an effective approach for targeting cranial NCCs, we next asked whether the system could achieve functional genome editing during craniofacial development. HH9/10 chicken embryos were injected with MC3 LNPs containing wild-type Cas9 mRNA and synthetic guide RNAs targeting *RUNX2*, a transcription factor with a central role in craniofacial skeletogenesis (Hall et al. 2014; Newton and Pask, 2020). To quantify the efficiency and spatial specificity of editing, embryos were collected four days after injection, and the frontonasal process (FNP) and combined mandibular/maxillary (MAN/MAX) prominences were dissected from the injected right and contralateral left sides. Targeted amplicon sequencing was used to quantify the proportion of edited *RUNX2* alleles.

Substantial *RUNX2* editing was detected specifically within the LNP-injected side of both craniofacial regions (Fig. 6A). Approximately 40% of *RUNX2* alleles were edited in the injected MAN/MAX complex, while editing approached 50% in the injected FNP. In contrast, editing remained low in the corresponding contralateral tissues and was negligible in uninjected controls. Because sequencing was performed on total dissected tissue rather than an enriched population of transfected cells, these editing frequencies demonstrate substantial genome modification at the tissue level following localized LNP delivery in ovo.

**Fig. 6.**
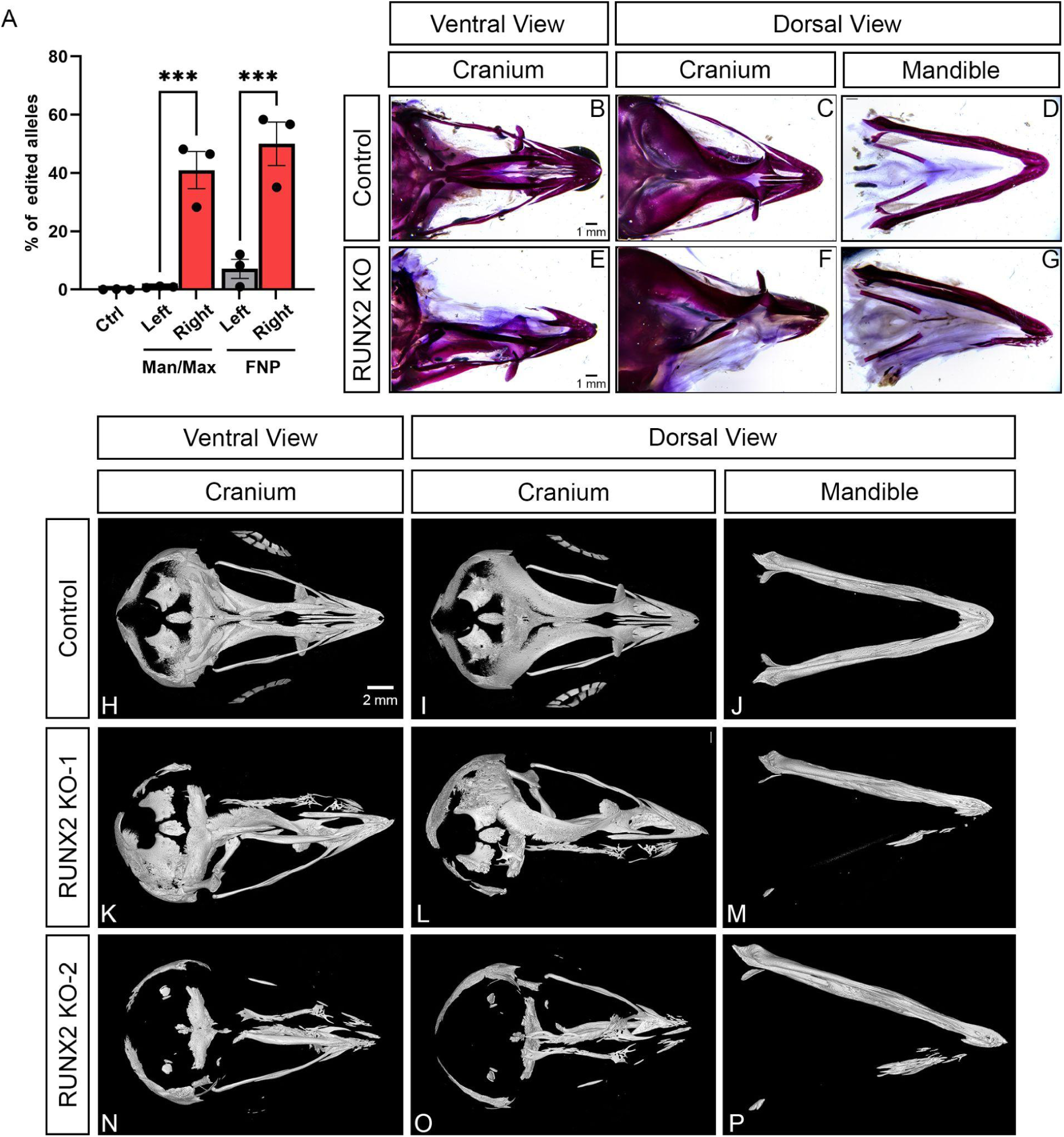
**LNP-mediated CRISPR/Cas9 delivery enables efficient RUNX2 genome editing and produces severe craniofacial skeletal defects in chicken embryos**. HH9/10 chicken embryos were injected with MC3 LNPs containing wild-type Cas9 mRNA and synthetic guide RNAs targeting *RUNX2* into the right cephalic mesenchyme adjacent to the neural tube. The left side served as an internal non-injected control. (A) Quantification of *RUNX2* editing in the combined mandibular/maxillary (MAN/MAX) prominences and frontonasal process (FNP). Tissues from the injected right and left sides were dissected 4 days after injection, dissociated and subjected to targeted Next Generation Sequencing (NGS) of amplicons. Editing efficiency is expressed as the percentage of *RUNX2* alleles containing edits. (B–G) Alizarin Red staining of craniofacial skeletal structures at HH42. Control embryo shown in ventral (B) or dorsal (C) view of the cranium, and dorsal view of the mandible (D). *RUNX2*-edited embryo shown in the corresponding ventral (E) or dorsal (F) view of the cranium, and dorsal view of the mandible (G). (H–P) Three-dimensional micro-CT analysis of craniofacial skeletal morphology at HH42. Control embryo shown in ventral view of the cranium (H), dorsal view of the cranium (I), and dorsal view of the mandible (J). Two independently edited embryos, RUNX2 KO-1 (K–M) and RUNX2 KO-2 (N–P) are shown in the corresponding ventral cranial (K, N), dorsal cranial (L, O) and mandibular (M, P) views. RUNX2-edited embryos exhibit severe loss and disruption of mineralized craniofacial skeletal structures. Scale bars: 1 mm (B–G) and 2 mm (H–P). Data in A are presented as mean ± s.e.m.; individual points represent independent samples. Statistical significance is indicated in the figure; ***P<0.001.

To determine whether this level of somatic editing was sufficient to alter craniofacial development, injected embryos were allowed to develop to HH42 and craniofacial skeletal morphology was assessed by Alizarin Red staining and micro-computed tomography (micro-CT). Compared with controls, RUNX2-edited embryos showed pronounced loss and disruption of mineralized craniofacial structures, affecting both the cranium and mandible (Fig. 6B–G). Three-dimensional micro-CT analysis confirmed extensive loss and malformation of cranial and mandibular skeletal elements in independently edited embryos (Fig. 6H–P). Thus, LNP-mediated delivery of CRISPR/Cas9 components at HH9/10 generated sufficient somatic *RUNX2* disruption to produce a severe craniofacial skeletal phenotype at later embryonic stages.

These results demonstrate that LNPs can achieve high levels of spatially localized somatic genome editing in avian embryos and, importantly, that the resulting editing is sufficient to produce a readily detectable developmental phenotype.

## DISCUSSION

Here, we establish lipid nanoparticles as a versatile platform for nucleic acid delivery and functional genetic manipulation in developing avian embryos. Localized LNP administration produced reproducible reporter expression within targeted craniofacial tissues while maintaining high short-term embryo survival, and repeated administration substantially increased cellular coverage. The same delivery framework accommodated multiple nucleic acid cargos and experimental applications, ranging from transient mRNA expression to persistent piggyBac-mediated labeling and CRISPR/Cas9 genome editing. Importantly, LNP-mediated delivery of Cas9 mRNA and synthetic guide RNAs generated substantial somatic editing of *RUNX2* and produced a pronounced craniofacial skeletal phenotype. Together, these findings demonstrate that LNPs can provide sufficient delivery efficiency for functional interrogation of gene activity directly within the developing avian embryo.

LNP delivery provides several features that complement existing approaches for genetic manipulation in avian embryos. Electroporation remains a powerful and widely used method, but transfection efficiency and tissue distribution can depend strongly on electrode positioning, tissue geometry, and electrical parameters (Simkin et al., 2014). Localized LNP microinjection does not require application of an electrical field and, under the conditions tested here, reproducibly targeted the developing cephalic mesenchyme. Because LNP delivery requires only microinjection access to the target tissue, this strategy may be particularly advantageous for extending in ovo genetic manipulation to species with smaller eggs, where the physical space required for conductive solution and electrode positioning can constrain electroporation. Moreover, the high survival rates of embryos treated with LNPs bodes well for studies of avian species where obtaining large numbers of eggs is a rate-limiting step.

The ability to administer LNPs repeatedly during development provides an additional advantage for studying dynamic cell populations. Sequential injections progressively increased the proportion of cranial NCCs receiving cargo while also allowing previously targeted cells to receive additional cargo. Thus, repeated administration can both access cells entering the target region over time and re-expose previously targeted populations, a feature that may be particularly useful for developmental processes involving cell migration or for applications requiring efficient delivery of multiple genetic components.

A major advantage of the platform is its flexibility with respect to nucleic acid cargo and the duration of genetic manipulation. LNP-mediated mRNA delivery enabled rapid transient reporter expression, whereas delivery of the piggyBac system supported persistent reporter labeling through advanced stages of craniofacial development. Although genomic integration was not directly assayed in this study, the persistence of reporter-positive cells within differentiated craniofacial derivatives is consistent with the expected activity of the piggyBac system and demonstrates the utility of LNP delivery for long-term genetic labeling. Most importantly, LNPs efficiently co-delivered Cas9 mRNA and synthetic guide RNAs, producing approximately 40–50% edited *RUNX2* alleles within dissected craniofacial tissues. Because these measurements were obtained from total tissue rather than an enriched population of transfected cells, they indicate substantial editing across the targeted region. This level of somatic editing, which targeted NCC on one side of the embryos, was sufficient to produce severe unilateral disruption of craniofacial ossification, demonstrating that LNP-mediated editing can target specific cell populations and generate developmental phenotypes without first establishing a stable genetically modified line.

The flexibility of LNP formulation provides opportunities for further optimization, but also creates a substantial parameter space as the approach is extended to additional applications. LNP performance can be influenced by ionizable lipid identity, helper lipid composition, component ratios, cargo-to-lipid ratios, and the properties of the cargo itself. In addition, localized delivery requires physical access to the target region, and the present study primarily evaluates cranial NCCs and their derivatives. Whether similar efficiencies can be achieved in more densely organized tissues or other embryonic cell populations remains to be determined. Defining how formulation, injection strategy, developmental stage, and target tissue influence delivery efficiency will therefore be important for establishing the broader applicability of the platform.

Finally, successful LNP-mediated delivery and persistent reporter labeling in chicken, pigeon, and duck embryos demonstrate that the approach is not restricted to a single avian species. This is particularly relevant for comparative developmental and evolutionary studies, where functional genetic experiments have historically been concentrated in a small number of experimentally tractable and commercially available models. Extending LNP-based delivery to additional avian species could provide a common experimental framework for directly testing the developmental consequences of genetic and regulatory variation across birds. More broadly, the combination of spatially localized delivery, repeat dosing, cargo flexibility, and efficient somatic genome editing provides a foundation for expanding functional genetic approaches in avian developmental biology and evolution.

## MATERIALS AND METHODS

### Animal welfare

All procedures involving avian embryos were performed in accordance with institutional guidelines, applicable federal regulations governing the use of avian embryos in research, and accepted practices for humane treatment as described in the AVMA Guidelines (Leary et al. 2020). Consistent with guidance from the U.S. National Institutes of Health Office of Laboratory Animal Welfare (OLAW), embryos of oviparous species are not considered live vertebrate animals prior to hatching. No embryos were allowed to hatch and chicken (*G. gallus*) and domestic pigeon (*C. livia*) embryos were incubated for up to 17 days, duck (*A. platyrhynchos*) embryos were incubated for up to 19 days.

### Embryo preparation and in ovo LNP delivery

Fertilized chicken (*G. gallus*), domestic pigeon (*C. livia*) and duck (*A. platyrhynchos*) eggs were incubated at 38°C and 60% relative humidity in a humidified incubator (Sportsman 1502, GQF Manufacturing, Savannah, GA, USA) until the desired developmental stage. To increase viability, pigeon and duck eggs were incubated with continuous rocking and placed in a horizontal position one to two hours before opening the egg to allow the embryo to position in the right orientation. Embryos were staged according to the Hamburger-Hamilton (HH) staging system (Hamburger and Hamilton, 1951; Hamilton, 1965). A 1–2 cm window was opened in the eggshell above the embryo using sterile scissors. To facilitate visualization, approximately 0.2–0.3 ml of filter-sterilized 0.1% India ink diluted in Hanks’ Balanced Salt Solution (HBSS; Thermo Fisher Scientific, Cat# 14-175-095) was injected beneath the embryo using a hypodermic syringe.

LNP formulations containing the indicated DNA or RNA cargo were delivered using custom-pulled aluminosilicate glass capillary needles (Sutter Instrument, Cat# AF100-64-10). Capillary needles were prepared using a P-1000 micropipette puller (Sutter Instrument) with the following parameters: (heat, 450; pull, 150; velocity, 150; time, 150; pressure, 150). LNP suspensions were supplemented with 0.1% Fast Green FCF (Sigma-Aldrich, Cat# F7252) to visualize injected material. Needles were backfilled with the LNP suspension using microloader tips (Calibre Scientific, Cat# EPE930001007). Embryos were injected at HH9/10 into the right cephalic mesenchyme immediately adjacent to the developing neural tube to target migrating cranial NCCs. During each injection session, LNP suspension was delivered across 3–5 adjacent positions within the cephalic mesenchyme, for a total injected volume of approximately 100–150 nL per session. Microinjections were performed using a pneumatic PicoPump (World Precision Instruments, Cat# SYS-PV830) at 3–5 PSI. Only the right side was injected; the contralateral left side served as an internal non-injected control. For repeated-delivery experiments, embryos received up to three injection sessions separated by 2–3 h. After each session, the eggshell window was sealed with transparent adhesive tape and embryos were returned to the incubator until the next injection or collection time point.

### Evaluation of transgene expression

Reporter expression was initially evaluated 24 hours after the first LNP injection by detection of EGFP or mCherry fluorescence. Embryos were imaged using a Leica THUNDER stereomicroscope (Leica Microsystems). Reporter expression was evaluated within the injected cephalic region and compared with the contralateral non-injected side of the same embryo where applicable.

### Craniofacial tissue collection

Following LNP injection, embryos were returned to the incubator and allowed to develop to the indicated HH stages. Embryos were collected in ice-cold phosphate-buffered saline (PBS), and craniofacial tissues were microdissected under a stereomicroscope. Depending on the experiment, the frontonasal prominence (FNP), maxillary prominence (MAX), and mandibular prominence (MAN) were collected individually or in the combinations indicated for downstream analysis. Dissected tissues were transferred immediately to pre-chilled tubes and processed for flow cytometry or next-generation sequencing (NGS), as appropriate.

### Embryo Viability

Embryo viability was assessed at the indicated time points by visual confirmation of cardiac activity and normal circulation. Embryos lacking detectable cardiac activity were scored as non-surviving. Survival was calculated as the number of viable embryos divided by the total number of embryos evaluated for each treatment group. Embryos lost because of pathogenic contamination or egg damage were excluded from the analysis.

### Next Generation Sequencing

Genomic DNA was extracted from individual tissue samples using QuickExtract™ DNA Extraction Solution (Biosearch Technologies, Cat# QE09050) according to the manufacturer’s protocol. Tissue was resuspended in 30–50 µL QuickExtract solution and incubated at 65°C for 6 min and 98°C for 2 min, followed by a 4°C hold. Genomic DNA was then used for targeted amplicon sequencing. PCR1 amplification was performed using KOD One™ PCR Master Mix Blue (Toyobo, Cat# KMM-201) under the following cycling conditions: 98°C for 30 s; 34 cycles of 98°C for 10 s, 60°C for 5 s, and 68°C for 5 s; 68°C for 30 s; hold at 4°C. PCR1 primers included universal adapter sequences for PCR2 amplification (F: 5′-TCGTCGGCAGCGTCAGATGTGTATAAGAGACAG-3′ and R: 5′-GTCTCGTGGGCTCGGAGATGTGTATAAGAGACAG-3′.) *RUNX2* primers were F: 5′-GCAAGATGAGCGAGGTCAG-3′ and R: 5′-GCTGTCAGCGCGGGGG-3′. PCR1 amplicons were indexed using xGen^TM^ UDI 10nt primer pairs (Integrated DNA Technologies, Cat# 10008054) under the following cycling conditions: 10 cycles of 98°C for 10 s, 60°C for 5 s, and 68°C for 1 s; 68°C for 10 s; hold at 4°C. Indexed amplicons were then pooled and cleaned using a 1:1 ratio of AmPure XP beads (Beckman Coulter, Cat# A63881). Library pools were sequenced on the MiSeq i100 (Illumina).

### Flow Cytometry

Individual tissue samples were processed into single-cell suspensions by incubation in 100-200 µL of 1X TrypLE™ Express Enzyme (Thermo Fisher Scientific, Cat# 12605010) at 37 °C for 30 min, with manual resuspension by pipetting every 10 min. Digestion was stopped by adding two volumes of DMEM containing 10% FBS. Cell suspensions were passed through 35 µm Falcon Cell Strainer Snap Caps (Corning, Cat# 352235), resuspended in Cell Staining Buffer (BioLegend, Cat# 420201), and stained for 20 min at room temperature with anti-human CD57 antibody (BioLegend, clone HNK-1, Cat# 359608). Samples were analyzed on an Attune NxT Flow Cytometer (Thermo Fisher Scientific), and data were analyzed using FlowJo v10.10.

### Cell culture

DF-1 chicken embryo fibroblasts (ATCC, CRL-3586) were cultured in Dulbecco’s Modified Eagle Medium (DMEM) supplemented with 10% fetal bovine serum (FBS), 1% penicillin-streptomycin, 1x GlutaMAX, 1x sodium pyruvate and maintained at 37°C in a humidified atmosphere containing 5% CO₂. Cells were grown as an adherent monolayer in tissue culture-treated flasks and passaged every 2–3 days at approximately 80–90% confluence. Cells were imaged and analyzed using an Incucyte SX5 Live-Cell Analysis System (Sartorius).

### Lipid Nanoparticle Formulation

#### Materials

All lipids were obtained from Cayman Chemical and supplied as crystalline solids or solutions in ethanol at the stock concentrations listed in Table 1. The ionizable lipids, helper lipids, sterol, and PEG-lipid, together with catalog numbers, CAS numbers, and the molecular weights used for N:P calculations, are given in Table 1.

**Table 1.** Lipid reagents (all Cayman Chemical).

| Reagent | Cat. no. | CAS no. | MW (g/mol) | Stock (mg/mL) |
| --- | --- | --- | --- | --- |
| DLin-MC3-DMA (MC3) | 34364 | 1224606-06-7 | 642.1 | 10 |
| DLin-KC2-DMA (KC2) | 34363 | 1190197-97-7 | 642.1 | 10 |
| SM102 | 33474 (custom) | 2089251-47-6 | 710.2 | 10 |
| LP-01 | 37278 (custom) | 1799316-64-5 | 852.29 | 10 |
| DSPC | 15100 | 816-94-4 | 790.15 | 19.75 |
| DOPE | 15091 | 4004-05-1 | 744.03 | 10 |
| Cholesterol | 9003100 | 57-88-5 | 386.66 | 9.67 |
| DMG-PEG(2000) | 33945 | 160743-62-4 | 2,509.2 | 10 |

##### Solvent and buffer

Molecular-biology-grade absolute ethanol (200 proof; Sigma-Aldrich, cat. no. E7023) was used as the organic solvent, and LipidLaunch™ LNP Formulation Buffer (50 mM acetate, pH 4.5; Cayman Chemical, cat. no. 400813) as the aqueous phase.

##### Nucleic acid cargoes

Reporter messenger RNAs were mCherry mRNA (Cap1, N1-methylpseudouridine [m1Ψ]; GenScript, cat. no. RP-A00041-0.2) and eGFP mRNA (Cap1, m1Ψ; GenScript, cat. no. RP-A00009-0.2), which were used interchangeably. The editor mRNA was Cas9-P2A-mScarlet mRNA (custom synthesis; TriLink Biotechnologies), modified with N1-methylpseudouridine, CleanCap M6 capping, and a 100-nt poly(A) tail (plasmid-encoded poly(A) track), purified by oligo-dT chromatography and supplied in 1 mM sodium citrate (pH 6.4). Single-guide RNAs targeting RUNX2 were obtained by custom synthesis (Integrated DNA Technologies). Two plasmids, p23 (PiggyBac transposon donor) and CMV-PB (PiggyBac transposase expression), were produced in house and purified using the Macherey-Nagel NucleoBond Xtra Midi kit (cat. no. 740410.50).

#### Lipid nanoparticle preparation

Lipid nanoparticles (LNPs) were prepared by hand-mixing separate aqueous and organic phases in 1.5 mL microcentrifuge tubes. A panel of four ionizable lipids — DLin-MC3-DMA (MC3), DLin-KC2-DMA (KC2), SM102, and LP-01 — was evaluated across three cargo classes: reporter mRNA, Cas9 editor mRNA with single-guide RNA (sgRNA), and plasmid DNA. All formulations were prepared by the common procedure described below; formulation-specific lipid compositions, N:P ratios, cargoes, volumes, and final nucleic acid concentrations are summarized in Table 2.

**Table 2.**
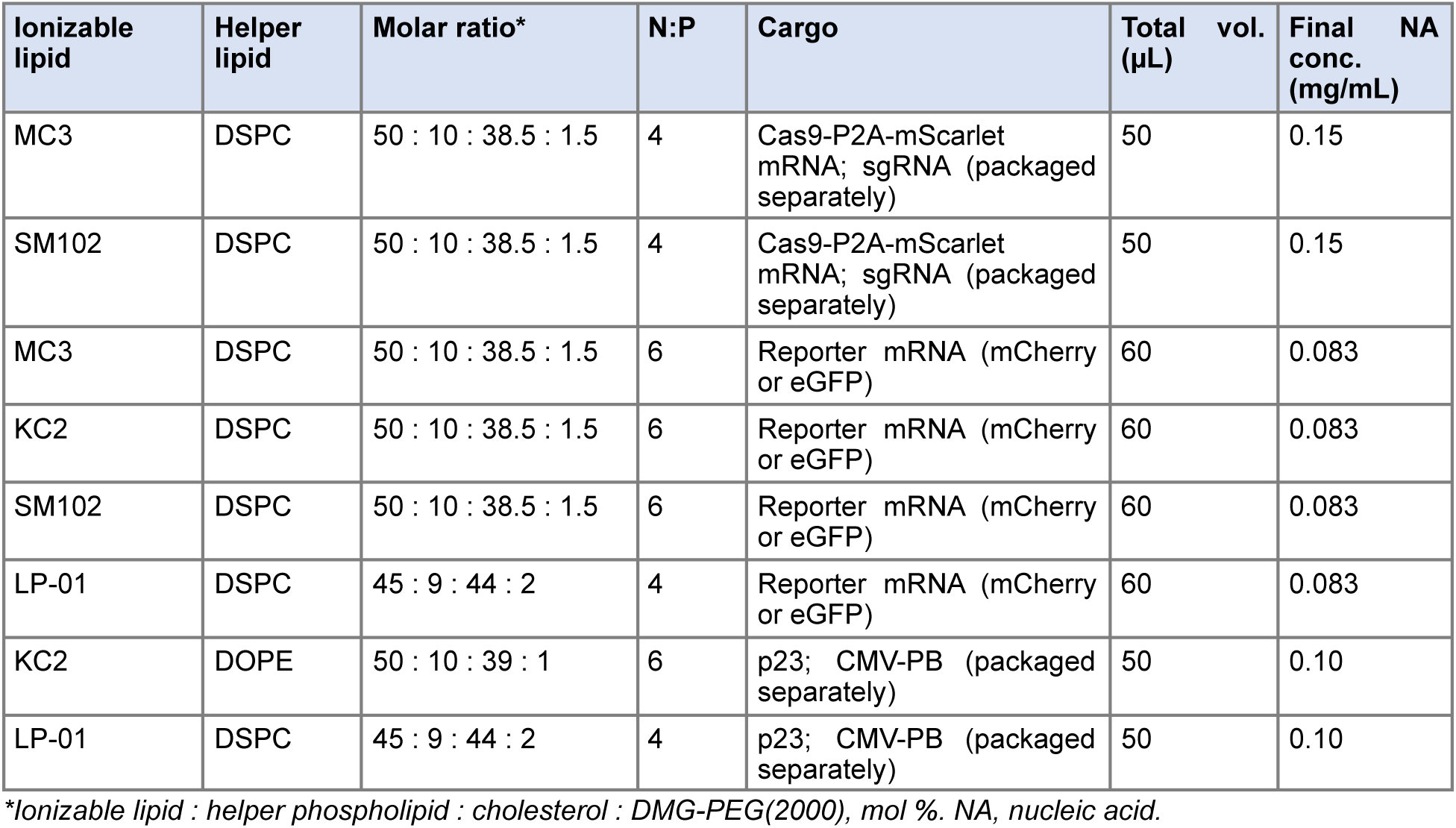
LNP formulations. Molar ratio is given as ionizable lipid : helper phospholipid : cholesterol : DMG-PEG(2000).

| Ionizable lipid | Helper lipid | Molar ratio* | N:P | Cargo | Total vol. ($\mu\text{L}$) | Final NA conc. (mg/mL) |
| --- | --- | --- | --- | --- | --- | --- |
| MC3 | DSPC | 50 : 10 : 38.5 : 1.5 | 4 | Cas9-P2A-mScarlet mRNA; sgRNA (packaged separately) | 50 | 0.15 |
| SM102 | DSPC | 50 : 10 : 38.5 : 1.5 | 4 | Cas9-P2A-mScarlet mRNA; sgRNA (packaged separately) | 50 | 0.15 |
| MC3 | DSPC | 50 : 10 : 38.5 : 1.5 | 6 | Reporter mRNA (mCherry or eGFP) | 60 | 0.083 |
| KC2 | DSPC | 50 : 10 : 38.5 : 1.5 | 6 | Reporter mRNA (mCherry or eGFP) | 60 | 0.083 |
| SM102 | DSPC | 50 : 10 : 38.5 : 1.5 | 6 | Reporter mRNA (mCherry or eGFP) | 60 | 0.083 |
| LP-01 | DSPC | 45 : 9 : 44 : 2 | 4 | Reporter mRNA (mCherry or eGFP) | 60 | 0.083 |
| KC2 | DOPE | 50 : 10 : 39 : 1 | 6 | p23; CMV-PB (packaged separately) | 50 | 0.10 |
| LP-01 | DSPC | 45 : 9 : 44 : 2 | 4 | p23; CMV-PB (packaged separately) | 50 | 0.10 |

\*Ionizable lipid : helper phospholipid : cholesterol : DMG-PEG(2000), mol %. NA, nucleic acid.

##### Organic phase

For each formulation, the ionizable lipid, a helper phospholipid (DSPC or DOPE), cholesterol, and DMG-PEG(2000) were combined in absolute ethanol at the molar ratio specified in Table 2, using the ethanolic stock solutions listed in Table 1; the combined volume was brought to the final organic-phase volume with absolute ethanol. The ionizable lipid content of each formulation was set to achieve the target N:P ratio (moles of ionizable amine to moles of nucleic acid phosphate; Table 2). To improve pipetting accuracy, organic lipid working mixes were prepared at 1.25× scale.

##### Aqueous phase

Nucleic acid cargo was diluted in LipidLaunch™ LNP Formulation Buffer (50 mM acetate, pH 4.5) to the aqueous-phase volume, corresponding to three-quarters of the total formulation volume (3:1 aqueous:organic). For reporter-mRNA LNPs, mCherry or eGFP mRNA (used interchangeably; 1.0 µg/µL stock) was formulated at 5 µg in 45 µL of aqueous phase per 60 µL preparation. For editor and guide LNPs, Cas9-P2A-mScarlet mRNA and the sgRNA were each formulated as separate LNP populations (7.5 µg RNA each) in 37.5 µL of aqueous phase per 50 µL preparation. For plasmid-DNA LNPs, the donor (p23; 3.148 µg/µL stock) and transposase (CMV-PB; 3.375 µg/µL stock) plasmids were likewise formulated as separate LNP mixes (5 µg DNA each) in 37.5 µL of aqueous phase per 50 µL preparation. Separately packaged LNP formulations— editor and guide RNAs, and p23 and CMV-PB — were combined at defined ratios at the point of administration.

#### Mixing and incubation

LNPs were formed by combining the aqueous and organic phases at a 3:1 (v/v) aqueous:organic ratio for the total volume given in Table 2. The mixture was combined by pipetting for 30 s, then incubated at room temperature for 15 min to allow for particle formation. No dilution, dialysis, or buffer-exchange step was performed. The resulting LNPs were used immediately or stored at 4 °C and used within 72 hours. All formulations can be scaled to the total reaction volume required for a given experiment — 50, 60, 100, and 200 µL preparations are used routinely — by adjusting the aqueous and organic phase volumes proportionally while holding the 3:1 aqueous:organic ratio, the lipid molar ratios, the N:P ratio, and the final nucleic acid concentration constant.

##### Particle characterization

Hydrodynamic size (Z-average diameter) and polydispersity index (PDI) were measured by dynamic light scattering, and zeta potential by electrophoretic light scattering, on a Malvern Panalytical Zetasizer Advance Ultra using the instrument’s protein analysis setting. For all measurements, LNPs were diluted 1 µL in 1000 µL of PBS and analyzed at 25 °C with a 120 s equilibration time, recording five measurements per sample. Size measurements were performed in polystyrene cuvettes (Sarstedt, cat. no. 67.754), and zeta-potential measurements used a Malvern Folded Capillary Zeta Cell (product code DTS1070).

##### Alizarin Red Skeletal Staining

Embryos were collected at the indicated developmental stages, and the heads were dissected. To facilitate removal of the skin, heads were immersed in hot water (65–70°C) for 20–30 s, after which the skin was carefully removed using forceps and spring scissors. Samples were fixed overnight in 95% ethanol with gentle rocking and subsequently transferred to 100% acetone and incubated overnight at room temperature for tissue defatting. Following acetone treatment, samples were washed in 70% ethanol for 6–8 hours with several changes of ethanol and then transferred to 1% potassium hydroxide (KOH) overnight. Embryo heads were stained by incubation in 0.1% Alizarin Red in 1% KOH for 4 h at room temperature with gentle rocking.

Samples were subsequently transferred to 1% KOH and gently rocked until the surrounding soft tissues became sufficiently cleared, typically for up to 1 week. Following clearing, samples were transferred through a graded glycerol series consisting of 20%, 50%, and 75% glycerol in 1% KOH, with incubation for at least 24 hours at each concentration. Samples were then transferred to 100% glycerol and maintained until the remaining soft tissue was transparent and only the Alizarin Red-stained mineralized skeletal elements were clearly visible. Depending on the developmental stage and degree of tissue clearing, the complete glycerol clearing process required up to 3 weeks. Stained specimens were maintained in 100% glycerol for imaging and subsequent analysis.

##### Micro-computed tomography

Craniofacial skeletal morphology was assessed by micro-computed tomography (micro-CT) using a Bruker Skyscan 1276, at 45 kV and 177 µA with no beam filter, over a 360° rotation, at 7.0 µm isotropic voxel size. Cross-sections were reconstructed in NRecon v2.1.0.1 (Bruker) with ring-artifact and 20% beam-hardening corrections and exported as 16-bit TIFF stacks. Volumes were loaded into 3D Slicer v5.12.4 (Fedorov et al., 2012) and renderings were generated using scalar opacity transfer functions with white bone on black background and identical thresholds set to the same linear attenuation coefficient for each volume. The cranium, mandible and hyloid were manually segmented and separated and images generated for the dorsal and ventral views of the cranium and mandible.

##### Image quantification

Image quantification for Figs. 2 and 3 was performed in FIJI (ImageJ v1.54p). Raw Leica .LOF files were imported using the Bio-Formats plugin. The total head area was manually outlined and measured from the corresponding bright-field image. Reporter-positive tissue was quantified by applying the mean intensity threshold to fluorescence images and measuring the area above threshold. Transfection coverage was calculated as the reporter-positive area divided by the total head area. Mean fluorescence intensity was measured as the mean gray value within the reporter-positive region. The same analysis workflow was applied consistently across experimental groups.

##### Statistical analysis

Statistical analyses were performed using GraphPad Prism v11.1.0 (GraphPad Software, Boston, MA, USA). Each embryo was considered an independent biological replicate, with one egg representing one experimental unit (*n*=1). Data are presented as mean ± s.e.m., unless otherwise indicated. Comparisons among experimental groups were performed using one-way or two-way analysis of variance (ANOVA), as appropriate for the experimental design, followed by Tukey’s multiple-comparisons test. Statistical significance was defined as P<0.05 and is indicated in the figures as follows: *P<0.05, **P<0.01, ***P<0.001, and ****P<0.0001.

## REFERENCES

1. Ealba, E. L., Jheon, A. H., Hall, J., Curantz, C., Butcher, K. D. and Schneider, R. A. (2015). Neural crest-mediated bone resorption is a determinant of species-specific jaw length. Developmental Biology 408, 151–163. doi: 10.1016/j.ydbio.2015.10.001.

2. Fedorov, A., Beichel, R., Kalpathy-Cramer, J., Finet, J., Fillion-Robin, J.-C., Pujol, S., Bauer, C., Jennings, D., Fennessy, F., Sonka, M. et al. (2012). 3D Slicer as an image computing platform for the Quantitative Imaging Network. Magn. Reson. Imaging 30, 1323–1341. doi:10.1016/j.mri.2012.05.001.

3. Finn, J. D., Smith, A. R., Patel, M. C., Shaw, L., Youniss, M. R., van Heteren, J., Dirstine, T., Ciullo, C., Lescarbeau, R., Seitzer, J., et al. (2018). A single administration of CRISPR/Cas9 lipid nanoparticles achieves robust and persistent in vivo genome editing. Cell Reports 22, 2227–2235. doi: 10.1016/j.celrep.2018.02.014.

4. Gillmore, J. D., Gane, E., Taubel, J., Kao, J., Fontana, M., Maitland, M. L., Seitzer, J., O’Connell, D., Walsh, K. R., Wood, K. et al. (2021). CRISPR-Cas9 in vivo gene editing for transthyretin amyloidosis. New England Journal of Medicine 385, 493–502. doi: 10.1056/NEJMoa2107454.

5. Hall, J., Jheon, A. H., Ealba, E. L., Eames, B. F., Butcher, K. D., Mak, S.-S., Ladher, R., Alliston, T. and Schneider, R. A. (2014). Evolution of a developmental mechanism: species-specific regulation of the cell cycle and the timing of events during craniofacial osteogenesis. Developmental Biology 385, 380–395. doi: 10.1016/j.ydbio.2013.11.011.

6. Hamburger, V. and Hamilton, H. L. (1951). A series of normal stages in the development of the chick embryo. Journal of Morphology 88, 49–92.

7. Hamilton, H. L. (1965). *Lillie’s Development of the Chick: An Introduction to Embryology*. New York: Holt, Rinehart and Winston.

8. Hou, X., Zaks, T., Langer, R. and Dong, Y. (2021). Lipid nanoparticles for mRNA delivery. Nature Reviews Materials 6, 1078–1094. doi: 10.1038/s41578-021-00358-0.

9. Hughes, S. H. (2004). The RCAS vector system. Folia Biologica (Praha) 50, 107–119. doi:10.14712/fb2004050030107.

10. Newton AH, Pask AJ. 2020. Evolution and expansion of the RUNX2 QA repeat corresponds with the emergence of vertebrate complexity. Commun Biol 3:771

11. Noden DM, Schneider RA. 2006. Neural crest cells and the community of plan for craniofacial development: historical debates and current perspectives. Adv Exp Med Biol 589:1–23

12. Schneider, R. A. (2024). Cellular, molecular, and genetic mechanisms of avian beak development and evolution. Annual Review of Genetics 58, 433–454. doi: 10.1146/annurev-genet-111523-101929.

13. Schneider, R. A. and Helms, J. A. (2003). The cellular and molecular origins of beak morphology. Science 299, 565–568. doi: 10.1126/science.1077827.

14. Simkin, J. E., Zhang, D., Ighaniyan, S. and Newgreen, D. F. (2014). Parameters affecting efficiency of in ovo electroporation of the avian neural tube and crest. Developmental Dynamics 243, 1440–1447. doi: 10.1002/dvdy.24163.

15. Véron, N., Qu, Z., Kipen, P. A. S., Hirst, C. E. and Marcelle, C. (2015). CRISPR mediated somatic cell genome engineering in the chicken. Developmental Biology 407, 68–74. doi: 10.1016/j.ydbio.2015.08.007.

